# Pre-clinical studies of a recombinant adenoviral mucosal vaccine to prevent SARS-CoV-2 infection

**DOI:** 10.1101/2020.09.04.283853

**Authors:** Anne C. Moore, Emery G. Dora, Nadine Peinovich, Kiersten P. Tucker, Karen Lin, Mario Cortese, Sean N. Tucker

**Author notes:** Corresponding author. Dr. Sean Tucker, Chief Scientist Officer, Vaxart, Inc.

## Abstract

There is an urgent need to develop efficacious vaccines against SARS-CoV-2 that also address the issues of deployment, equitable access, and vaccine acceptance. Ideally, the vaccine would prevent virus infection and transmission as well as preventing COVID-19 disease. We previously developed an oral adenovirus-based vaccine technology that induces both mucosal and systemic immunity in humans. Here we investigate the immunogenicity of a range of candidate adenovirus-based vaccines, expressing full or partial sequences of the spike and nucleocapsid proteins, in mice. We demonstrate that, compared to expression of the S1 domain or a stabilized spike antigen, the full length, wild-type spike antigen induces significantly higher neutralizing antibodies in the periphery and in the lungs, when the vaccine is administered mucosally. Antigen-specific CD4+ and CD8+ T cells were induced by this leading vaccine candidate at low and high doses. This fulllength spike antigen plus nucleocapsid adenovirus construct has been prioritized for further clinical development.

## INTRODUCTION

The emergence of a novel coronavirus, severe acute respiratory syndrome coronavirus 2 (SARS-CoV-2), the causative agent of COVID-19 disease, in 2019, has led to a global pandemic and significant morbidity, mortality and socio-economic disruption not seen in a century. Coronavirus disease 2019 (COVID-19) is a respiratory illness of variably severity; ranging from asymptomatic infection to mild infection, with fever and cough to severe pneumonia and acute respiratory distress^1^. Current reports suggest that asymptomatic spread is substantial^2^, and SARS-CoV-2 infection induces a transient antibody response in most individuals^3^. Therefore, development of successful interventions is an immediate requirement to protect the global population against infection and transmission of this virus and its associated clinical and societal consequences. Mass immunization with efficacious vaccines has been highly successful to prevent the spread of many other infectious diseases and can also prevent disease in the vulnerable through the induction of herd immunity. Significant effort and resources are being invested in urgently identifying efficacious SARS-CoV-2 vaccines. A number of different vaccine platforms have demonstrated pre-clinical immunogenicity and efficacy against pneumonia^4, 5^. Several vaccines have demonstrated phase I or phase II safety and immunogenicity^6-8^. However, at this time, no vaccine has demonstrated efficacy in the field.

The most advanced SARS-CoV-2 vaccine candidates are all given by the intramuscular (IM) route, with some requiring −80 °C storage. This is a major barrier for vaccine dissemination and deployment during a pandemic in which people are asked to practice social distancing and avoid congregation. The ultimate goal of any vaccine campaign is to protect against disease by providing enough herd immunity to inhibit viral spread, not to make a set number of doses of vaccine. An injected solution takes a long period of time to administer and distribute and requires costly logistics, which means dose availability does not immediately translate to immunity. Further, systemic immunization can induce immunity in the periphery and lower respiratory tract. However, these vaccines cannot induce mucosal immunity in the upper respiratory tract. Mucosal IgA (with the polymeric structure and addition of the secretory component), creates more potent viral neutralization^9^, can block viral transmission^10, 11^, and in general, is more likely to create sterilizing immunity given that this is the first line of defense for a respiratory pathogen.

Mucosal vaccines can induce mucosal immune responses, antibodies and T cells at wet surfaces. We are developing oral vaccines for multiple indications, including influenza and noroviruses, delivered in a tablet form for people. Our vaccine platform is a replication-defective adenovirus type-5 vectored vaccine that expresses antigen along with a novel toll-like receptor 3 agonist as an adjuvant. These vaccines have been well tolerated, and able to generate robust humoral and cellular immune responses to the expressed antigens^12-14^. Protective efficacy in humans was demonstrated against a respiratory virus 90 days or more post vaccination, as shown in a well characterized experimental influenza infection model^15^. Furthermore, the vaccine also has the advantage of room temperature stability and needle-free, ease of administration, providing several advantages over injected vaccine approaches with respect to vaccine deployment and access.

Here, we describe the pre-clinical development of a SARS-CoV-2 vaccine based on Vaxart’s oral adenovirus platform. The key approach was to develop several vaccine candidates in parallel, in order to create premanufacturing seeds while initial immunogenicity experiments were in progress. Given that the vaccines were made during the pandemic, rapid decisions were required to keep the manufacturing and regulatory timelines from slipping. We assessed the relative immunogenicity of four candidate vaccines that expressed antigens based on the spike (S) and nucleocapsid (N) SARS-CoV-2 proteins. These proteins have been well characterized as antigens for related coronaviruses, such as SARS-CoV and MERS (reviewed in Yong, *et al*.,^16^) and, increasingly, for SARS-CoV-2 spike. The aim of our vaccine is to induce immunogenicity on three levels; firstly, to induce potent serum neutralizing antibodies to S, secondly to induce mucosal immune responses, and thirdly to induce T cell responses to both vaccine antigens. This three-fold approach aims to induce robust and broad immunity capable of protecting the individual from virus infection as well as disease, promote rapid dissemination of vaccine during a pandemic, and to protect the population from virus transmission through herd immunity.

Here, we report the induction of neutralizing antibody (Nab), IgG and IgA antibody responses, and T cell responses in mice following immunization of rAd vectors expressing one or more SARS-CoV-2 antigens.

## RESULTS

### Vector Construction

Initially, three different rAd vectors were constructed to express different SARS-CoV-2 antigens. These were a vector expressing the full-length S protein (**rAd-S**), a vector expressing the S protein and the N protein (**rAd-S-N**), and a vector expressing a fusion protein of the S1 domain with the N protein (**rAd-S1-N**). The N protein of rAd-S-N was expressed under control of the human beta actin promoter, which is much more potent in human cells than mouse cells. An additional construct where the expressed S protein was fixed in a prefusion conformation (**rAd-S(fixed)-N**) was constructed at a later date as a control for exploring neutralizing antibody responses. These are described in Figure 1. Expression of the various transgenes was confirmed following infection of 293 cells using flow cytometry and monoclonal antibodies to the S or N protein (supplemental figure 1).

**Figure 1.**
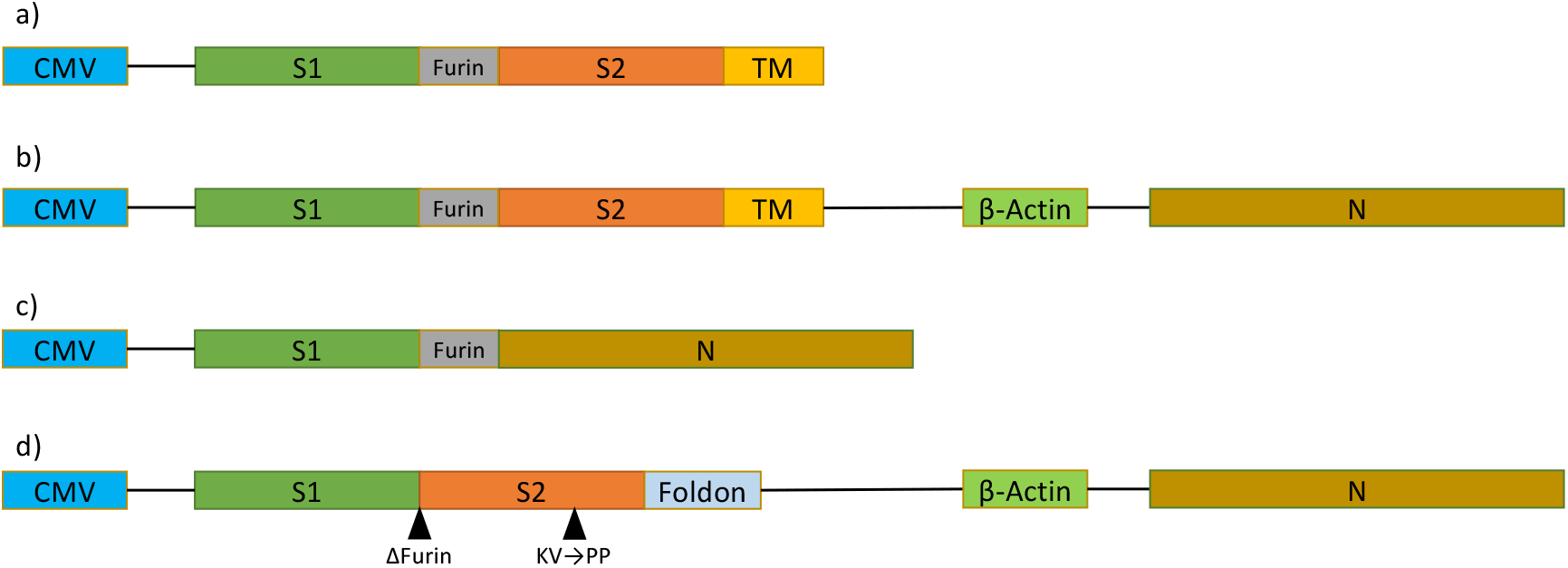
Transgene inserts developed to test vaccine specific responses. Recombinant adenoviruses were made using these inserts a. rAd-S b. rAd-S-N c. rAd-S1-N d. rAd-S(fixed)-N

### Immunogenicity of rAd vectors expressing S and N antigens

The primary objective of the initial mouse immunogenicity studies was to determine which of the rAd vectors induced significant antibody responses to S, and to obtain those results rapidly enough to provide a GMP seed in time for manufacturing. We and others^17^ have observed that transgene expression by vaccine vectors orally administered to mice can be suppressed in their intestinal environment, so immunogenicity was assessed following intranasal (i.n.) immunization. Animals were immunized i.n. and the antibody titers were measured over time by IgG ELISA. All three rAd vectors induced nearly equivalent anti-S1 IgG titers, at weeks 2 and 4 and the IgG titer in all animals was significantly boosted by the second immunization (p < 0.05 Mann Whitney t-tests) (Fig. 2A). However, the vector expressing full-length S (rAd-S-N) induced higher neutralizing titers compared to the vector expressing only S1 (Fig 2B). This was measured by two different neutralizing assays, one based on SARS-CoV-2 infection of Vero cells (cVNT) and one based on a surrogate neutralizing assay (sVNT). Furthermore, rAd-S-N induced higher lung IgA responses to S1 and unsurprisingly, to S2 (Fig. 2C) compared to rAd-S1-N two weeks after the final immunization. Notably, neutralizing titers in the lung were also significantly higher when rAd-S-N was used compared to the S1-containing vaccine (rAd-S1-N) (Fig. 2D). This demonstrated that the rAd-S-N candidate induced greater functional responses (NAb and IgA) compared to the vaccine containing the just the S1 domain. Because the N protein is much more highly conserved than the S protein, and is a target of long term T cell responses induced by infection^18^, the vector rAd-S-N was chosen for GMP manufacturing.

**Figure 2.**
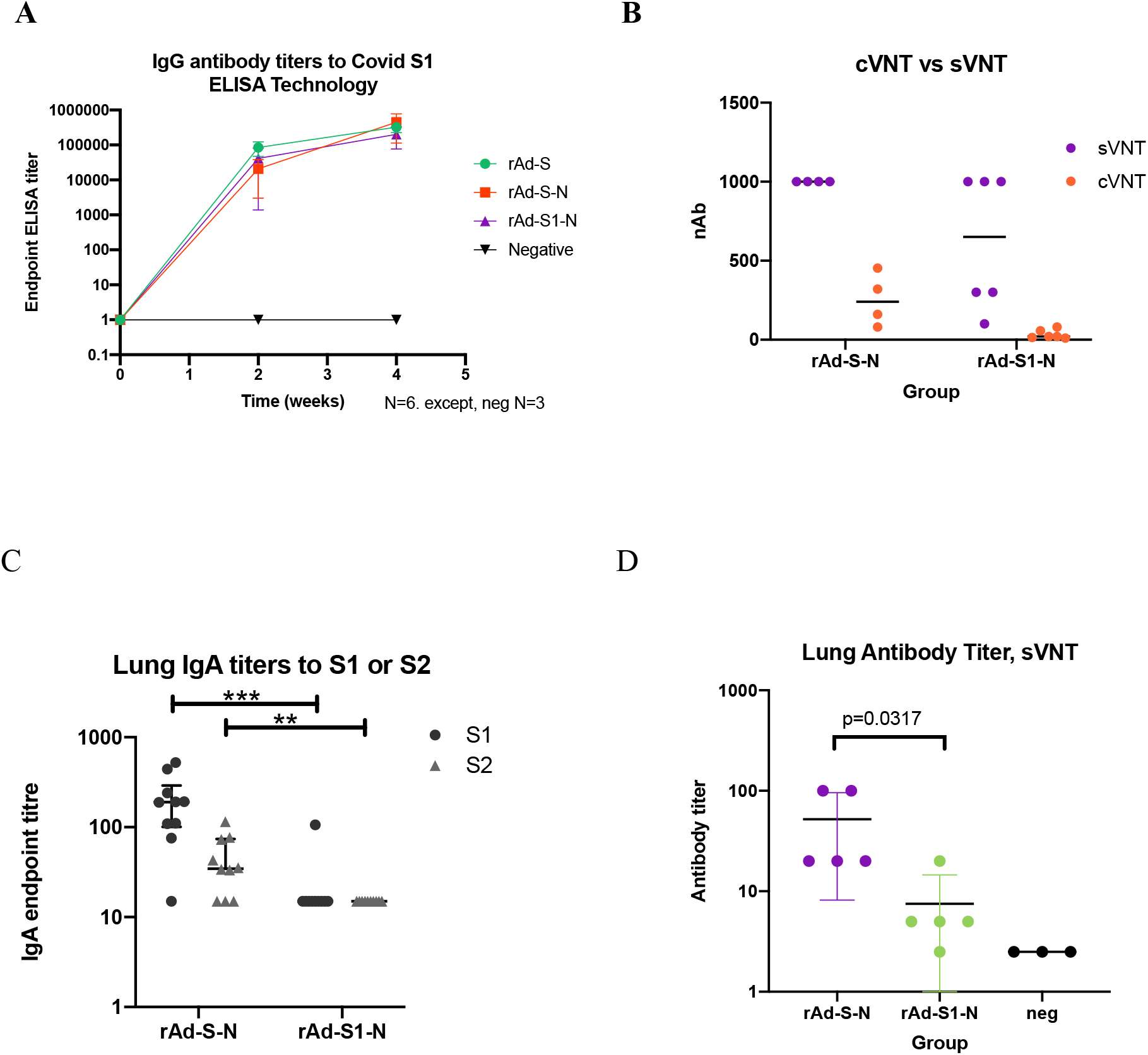
Immunization with candidate rAd vaccines induce serum IgG and lung IgA responses. Antibody titers to S following immunization of Balb/c mice on days 0 and 14 with 1x 10^8^ IU rAd expressing full-length S (rAd-S), co-expressing full length S and N (rAd-S-N) or co-expressing a fusion protein comprising the S1 domain and N (rAd-S1-N). **(A)** IgG serum IgG endpoint titers to S1 were measured by standard ELISA (n= 6 per vaccinated group, n=3 for PBS administered group). Symbols represent mean titers and bars represent the standard error) (**B)** Neutralizing antibody responses comparing rAd-S-N and rAd-S1-N using two different methods, surrogate VNT (sVNT) and cell-based VNT (cVNT). (**C**) IgA lung antibody titers to S1 and S2 in immunized mice. Endpoint titers were measured by standard ELISA (n= 10 per group). Lines represent the median and inter-quartile range. ** p<0.01, *** p < 0.001 defined by Mann-Whitney t-test. D. Neutralizing antibodies measured in the lungs post immunization.

Three dose levels of rAd-S-N were then tested to understand the dose responsiveness of this vaccine. The antibody responses to both S1 (Fig. 3A) and S2 (Fig. 3B) were measured. Similar responses were seen at all three dose levels at all timepoints. Responses to S1 and S2 were significantly increased at week 6 compared to earlier times, in all groups.

**Figure 3.**
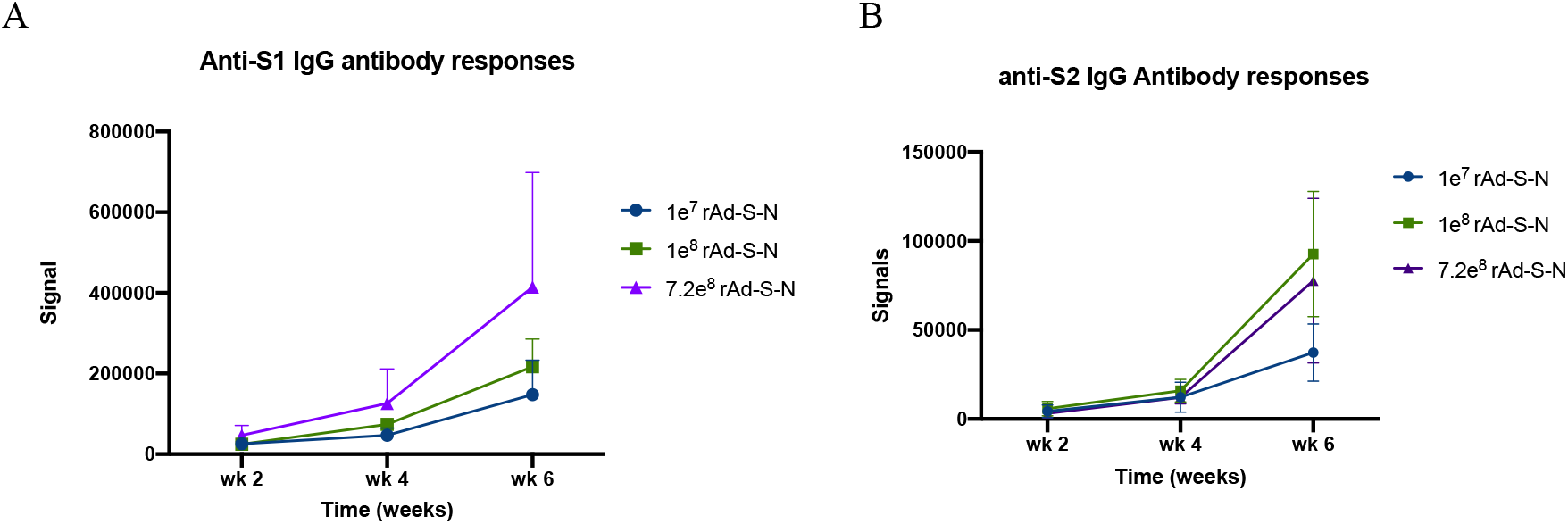
Immunization with rAd co-expressing full length S and N vaccines induce IgG responses in a dosedependent manner. **A and B.** Balb/c mice were immunized, IN, on days 0 and 14 with 1 x 10^7^ IU, 1 x 10^8^ IU or 7.2 x 10^8^ IU of rAd co-expressing full length S and N (rAd-S-N). The amount of IgG specific for S1 **(A)** and S2 in serum diluted 1/4000, was evaluated using a Mesoscale binding assay. Points represent the mean and lines represent the standard deviation.

The induction of S-specific T cells by rAd-S-N at different doses was then assessed. Induction of antigen-specific CD4+ and CD8+ T cells that produced effector cytokines such as IFN-γ, TNF-α and IL-2 was observed two weeks after 2 immunizations (Fig 4A). Notably, little IL-4 was induced by this vaccine and only in CD4+ T cells; providing a level of assurance that the risk for vaccine dependent enhancement of disease was very low. Furthermore, immunization with rAd-S-N induced double and triple positive, multi-functional IFN-γ, TNF-α and IL-2 CD4+ T cells (Fig. 4B). A second dose response experiment was performed to focus on T cell responses to the S protein, 4 weeks after the final immunization (week 8 of the study). Splenocytes were stimulated overnight with a peptide library to the S protein, divided in two separate peptide pools. T cell responses in the two pools were summed and plotted (Fig. 4C). Animals administered the 1e7 IU and the 1e8 IU dose levels had significantly higher T cell responses compared to the untreated animals but produced a similar number of IFN-γ secreting cells to each other, demonstrating a dose plateau at the 1e7 IU dose. Notably, this T cell analysis was conducted 4 weeks after the second immunization, potentially after the peak of T cell responses.

**Figure 4.**
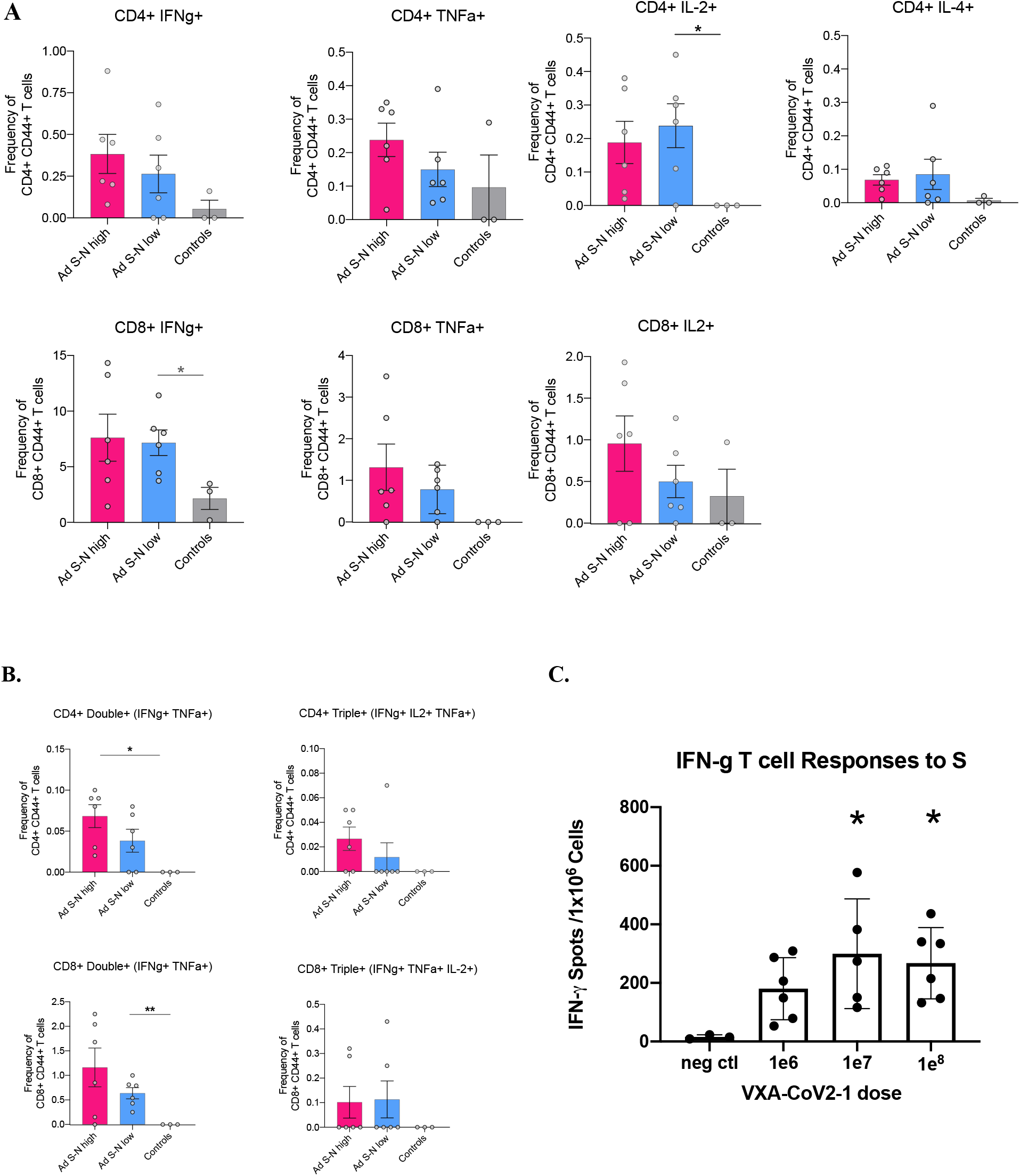
Immunization with rAd co-expressing full length S and N vaccines induce polyfunctional T cell responses. **(A)** Balb/c mice were immunized, IN, on days 0 and 14 with 1 x 10^8^ IU (Ad-S-Nhigh), 1 x 10^7^ IU (Ad-S-N low) of rAd-S-N. The frequency of CD4+ (top panel) or CD8+ T cells (bottom panel) that produced only IFN-γ, TNF-α, IL-2 or IL-4 after stimulation of spleen cells with 1μg/ml (CD4^+^) or 5μg/ml (CD8^+^) of the S peptide pools, as determined by ICS-FACS. **(B)** The frequency of polyfunctional CD4+ (top panel) or CD8+ T cells (bottom panel) that produced more than one cytokine after stimulation of spleen cells, Bars represent the mean and the lines represent the standard error of the mean. **(C)** IFN-γ T cell responses to S protein 4 weeks following immunization on weeks 0 and 4 with 1 x 10^6^ IU, 1 x 10^7^ IU, 1 x 10^8^ IU doses of rAd-S-N were measured by ELISPOT. Bars represent the mean and the lines represent the standard deviation. *p< 0.05; one-way non-parametric ANOVA with multiple comparisons.

### rAd-expressed wild-type S induces a superior neutralizing response compared to stabilized/pre-fusion S

An additional study was performed to compare rAd-S-N to a vaccine candidate with the S-protein stabilized and with the transmembrane region removed (rAd-S(fixed)-N). A stabilized version of the S protein has been proposed as a way to improve neutralizing antibody responses and produce less non-neutralizing antibodies. The S protein was stabilized through modifications as described by Amanat *et al*.,^19^. rAd-S-N induced higher serum IgG titers to S1 (Fig. 5A) at both timepoints tested, although these were not statistically significant at week 6 by Mann-Whitney (p = 0.067). However, rAd-S-N induced significantly higher neutralizing antibody responses (Fig. 5B) than the stabilized version (p = 0.0152). These results suggest that a wild-type version of the S protein is superior for a rAd based vaccine in mice.

**Figure 5:**
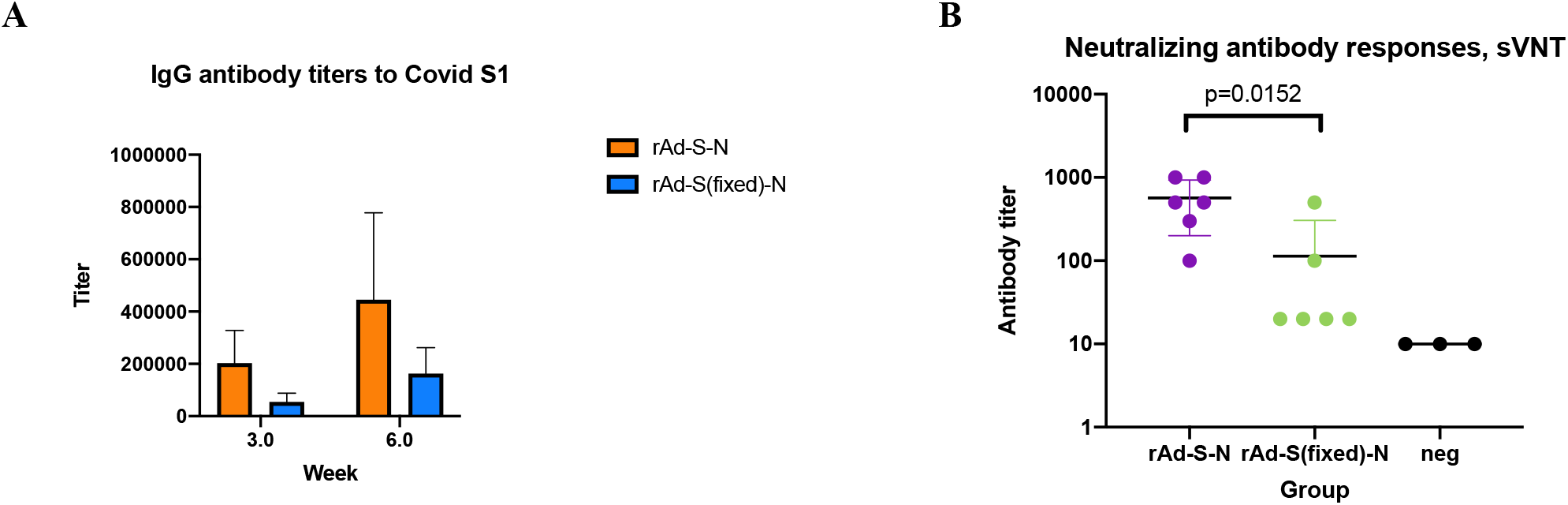
Antibodies to S were superior when the S protein expressed in the wild-type configuration compared to the fixed version. Balb/c mice were immunized on weeks 0 and 4 with 1e8 IU per mouse (n = 6), and antibody titers were measured. **(A)** IgG antibody titers over time. **(B**) Neutralizing antibody responses were measured at week 6. Note that 1:1000 was the maximum dilution performed.

## DISCUSSION

The endgame to the COVID-19 pandemic requires the identification and manufacture of a safe and effective vaccine and a subsequent global immunization campaign. A number of vaccine candidates have accelerated to phase III global efficacy testing and, if sufficiently successful in these trials, may form the first generation of an immunization campaign. However, all of these advanced candidates are S-based vaccines that are injected. Such an approach will unlikely prevent virus transmission, but should prevent pneumonia and virus growth and damage in the lower respiratory tract and periphery, as evidenced in macaque challenge studies^4, 5^.

One key constraint in a global COVID-19 immunization campaign will be the cold chain distribution logistics and a bottleneck of requiring suitably trained health care workers (HCWs) to inject the vaccine. Current logistics costs, including cold chain and training, can double the cost of fully immunizing an individual in a low-middle income country (LMIC)^20^. Implementing a mass immunization campaign, requiring trained HCWs for injection-based vaccines, will have a significant impact on healthcare resources in all countries. The need for cold chain, biohazardous sharps waste disposal and training will result in increased cost, inequitable vaccine access, delayed vaccine uptake and prolongation of this pandemic. These costs will be magnified if vaccines are unable to provide long-term protection (natural immunity to other beta-coronaviruses is short-lived^21^), and annual injection-based campaigns are needed. Vaxart’s oral tablet vaccine platform provides a solution to these immunological as well as logistic, economic, access and acceptability problems. In this study we demonstrate, in an animal pre-clinical model, the immunogenicity of a SARS-CoV-2 vaccine using Vaxart’s vaccine platform; namely the induction of serum and mucosal neutralising antibodies and poly-functional T cells.

Mouse studies were designed to test immunogenicity of candidate vaccines rapidly in the spring of 2020, before moving onto manufacturing and clinical studies critical to addressing the pandemic. Vaxart’s oral tablet vaccine platform has previously proven to be able to create reliable mucosal (respiratory and intestinal), T cell, and antibody responses against several different pathogens in humans^12, 14, 22, 23^. We know from our prior human influenza virus challenge study that oral immunization was able to induce protective efficacy 90 days post immunization; on par with the commercial quadrivalent inactivated vaccine^15^. These features provide confidence that the adoption of the platform to COVID-19 could translate to efficacy against this pathogenic coronavirus and could provide durable protection against virus infection. Finally, a tablet vaccine campaign is much easier because qualified medical support is not needed to administer it. This ease of administration will result in increased vaccine access and potentially, acceptability, as has been evidenced by the success of easy-to-administer, oral polio vaccine, in the elimination of polio virus^24^. These features could be even more important during SARS-CoV-2 immunization campaigns compared to other vaccines, as substantially more resources may be required to ensure uptake of this vaccine, given the global levels of COVID-19 denialism, mistrust and increased vaccine hesitancy^25, 26^. The tablet vaccine does not need refrigerators or freezers, does not require needles or vials, and can potentially be shipped via standard mail or by a delivery drone. These attributes significantly enhance deployment and distribution logistics, even permitting access to isolated regions with fewer technical resources. Finally, from an immunological perspective, oral administration of this adenovirus is not compromised by pre-existing immunity to adenoviruses nor creates substantial anti-vector immunity^12, 13^; issues that have been shown to cause significantly decreased vaccine potency in an rAd5 based SARS-CoV2 vaccine^27^ and can prevent durable increased immunity when the same adenovirus platform is re-administered by the IM route^28^.

The choice of antigen can be difficult during a novel pandemic, a time in which key decisions are needed quickly. The S protein is believed to be the major neutralizing antibody target for coronavirus vaccines, as the protein is responsible for receptor binding, membrane fusion, and tissue tropism. When comparing SARS-CoV-2 Wu-1 to SARS-CoV, the S protein was found to have 76.2% identity^29^. Both SARS-CoV and SARS-CoV-2 are believed to use the same receptor for cell entry: the angiotensin-converting enzyme 2 receptor (ACE2), which is expressed on some human cell types^30^. Thus, SARS–CoV-2 S protein is being used as the leading target antigen in vaccine development so far and is an ideal target given that it functions as the key mechanism for viral binding to target cells. However, the overall reliance on the S protein and an IgG serum response in the vaccines could eventually lead to viral escape. For influenza, small changes in the hemagglutinin binding protein, including a single glycosylation site, can greatly affect the ability of injected vaccines to protect^31^. SARS-CoV-2 appears to be more stable than most RNA viruses, but S protein mutations have already been observed without the selective pressure of a widely distributed vaccine. Once vaccine pressure begins, escape mutations might emerge. We took two approaches to address this issue; firstly to include the more conserved N protein in the vaccine and secondly to induce broader immune responses, namely through mucosal IgA.

High expression levels of ACE2 are present in type II alveolar cells of the lungs, absorptive enterocytes of the ileum and colon, and possibly even in oral tissues such as the tongue^32^. Transmission of the virus is believed to occur primarily through respiratory droplets and fomites between unprotected individuals in close contact^33^, although there is some evidence of transmission via the oral-fecal route as seen with both SARS-CoV and MERS-CoV viruses where coronaviruses can be secreted in fecal samples from infected humans^34^. There is also evidence that a subset of individuals exist that have gastrointestinal symptoms, rather than respiratory symptoms, are more likely to shed virus longer^35^. Driving immune mucosal immune responses to S at both the respiratory and the intestinal tract may be able provide broader immunity and a greater ability to block transmission, than simply targeting one mucosal site alone. Blocking transmission, rather than just disease, will be essential to reducing infection rates and eventually eradicating SARS-CoV-2. We have previously demonstrated that an oral, tableted rAd-based vaccine can induce protection against respiratory infection and shedding following influenza virus challenge^15^ as well as intestinal immunity to norovirus antigens in humans^12^. Furthermore, mucosal IgA is more likely to be able address any heterogeneity of the S proteins in circulating viruses than a monomeric IgG response. mIgA has also been found to be more potent at cross reactivity than IgG for other respiratory pathogens^36^. IgA may also be a more neutralizing isotype than IgG in COVID-19 infection, and in fact neutralizing IgA dominates the early immune response^37^. Notably, we saw a higher ratio between neutralizing to non-neutralizing antibodies in our lung versus serum antibody results in our mouse study as well, which supports the concept that IgA may have more potency compared to IgG. Polymeric IgA, through multiple binding interactions to the antigen and to Fc receptors can turn a weak single interaction into a higher overall affinity binding and activation signal, creating more cross-protection against heterologous viruses ^38^.

Our second strategy to mitigate this potential vaccine-driven escape problem was to include the N protein in the vaccine construct. The N protein is highly conserved among β-coronaviruses, (greater than 90% identical) contains several immunodominant T cell epitopes, and long-term memory to N can be found in SARS-CoV recovered subjects as well as people with no known exposure to SARS-CoV or SARS-CoV-2^18, 39^. In an infection setting, T cell responses to the N protein seem to correlate to increased neutralizing antibody responses^40^. All of these reasons led us to add N to our vaccine approach. The protein was expressed in 293A cells (Supp Fig1). However, as the human beta actin promoter is more active in human cells than mice, we did not explore immune responses in Balb/c mice, but will examine them more carefully in future NHP and human studies.

The optimum sequence and structure of the S protein to be included in a SARS-CoV-2 vaccine is a subject of debate. Several labs have suggested that reducing the S protein to the key neutralizing domains within the receptor binding domain (RBD) would promote higher neutralizing antibody responses, and fewer non-neutralizing antibodies^41, 42^. We made a vaccine candidate composed of the S1 domain, which includes the RBD, in an attempt to promote this approach. Although the S1-based vaccine produced similar IgG binding titers to S1, neutralizing antibody responses were significantly lower compared to the full-length S antigen. Other gene-based vaccines have also shown the reductionist approach to S does not work so well, demonstrating that the DNA vaccine expressing the full-length S-protein produced higher neutralizing antibodies than shorter S segments^5^. In agreement with these macaque studies, we observed that the sequence of the adenovirus encoded antigen had a significant impact on antibody function, here with respect to neutralization. While reducing the potential for exposing non-neutralizing antibody epitopes seems reasonable in theory, this might reduce the T cell help that allows for greater neutralizing antibodies to develop. Indeed, of the spike protein T cell responses, which make up 54% of the responses to SARS-CoV-2, only 11% map to receptor binding domain^43^. Stabilizing the S protein might be important for a protein vaccine, but not necessarily for a gene-based vaccine. The former is produced *in vitro* and it is produced to retain a homogenous, defined structure, ready for injection. In contrast, the latter, is expressed on the surface of a cell, *in vivo*, like natural infection, substantially in a prefusion form, and the additional stabilization may be unnecessary for B cells to create antibodies against the key neutralizing epitopes. We directly compared a stabilized version of S to the wild-type version. The wild-type version was significantly better at inducing neutralizing antibody responses. Of interest, this was also observed in a DNA vaccine study in NHPs, where the stabilized version appeared to induce lower neutralizing antibody (NAb) titers compared to the wild-type S^*5*^. A slightly different result was observed in studies of rAd26 vectors by Mercado, *et al* in NHPs, where expressing a stabilized version of the S protein appeared to improve NAb but lower T cell responses ^44^. In summary, stabilization doesn’t universally improve the immune responses in gene-based or vector-based vaccines.

Multiple vaccine candidates are in, or are about to begin, clinical testing. Due to known safety and immunogenicity for epidemic pathogens such as Ebola virus, two leading candidate vaccines are based on recombinant adenovirus vectors; University of Oxford’s ChAdOx1-nCov and Janssen Pharmaceutical’s AdVac platforms^45-48^. We saw stronger serum IgG and NAb titers in our study compared to a ChAdOx1-nCov in Balb/c mice^4^, however, this might reflect differences in assay components. A rAd36 vaccine study was performed by Hassan, *et al*., where doses of 1e10 VP were given by intranasal delivery^49^. The results were significant from the standpoint of blocking lung infection in a mouse SARS-CoV-2 challenge model. They reported titers of serum antibody titers of 1e4 above the background titers, similar to our results, despite using doses 2- to 3-log fold higher viral doses compared to our study. Indeed, in our study, equivalently strong T cell and antibody responses were observed using 1e7 IU and 1e8 IU by the intranasal route. Using these doses, we observed high percentages of CD8+ T cell responses (up to 14%) secreting IFN-γ and TNF-α and strong CD4+ T cells after peptide restimulation. Although we did not evaluate the trafficking properties of these antigen-specific T cells, we know that oral administration of this Ad-based vaccine in humans induces high levels of mucosal homing lymphocytes^12, 15^. A proportion of the antigenspecific CD4+ and CD8+ T cells were polyfunctional in this mouse study. Vaccine-induced T cells possessing multiple functions may provide more effective elimination of virus subsequent to infection and therefore could be involved in the prevention of disease. However, it is uncertain at this time what is the optimum T cell phenotype required for protection against disease.

In summary, these studies in mice represent our first step in creating a vaccine candidate, demonstrating the immunogenicity of the construct at even low vaccine doses and the elucidation of the full-length spike protein as a leading candidate antigen to induce T cell responses and superior systemic and mucosal neutralizing antibody. Future work will focus on the immune responses in humans.

## METHODS

### Vaccine Constructs

For this study, four recombinant adenoviral vaccine constructs were created based on the published DNA sequence of SARS-CoV-2 publicly available as Genbank Accession No. MN908947.3. Specifically, the published amino acid sequences of the SARS-CoV-2 spike protein (S protein) and the SARS-CoV-2 Nucleocapsid protein (N protein) were used to synthesize nucleic acid sequences codon optimized for expression in *Homo sapiens* cells (Blue Heron Biotechnology, Bothell, WA). These sequences were used to create recombinant plasmids containing transgenes cloned into the E1 region of Adenovirus Type 5 (rAd5), as described by He, *et al* ^50^, using the same vector backbone used in prior clinical trials for oral rAd tablets ^12, 15^. As shown in Fig 1, the following four constructs were created:

a. rAd-S: rAd5 vector containing full-length SARS-CoV-2 S gene under control of the CMV promoter.
b. rAd-S-N: rAd5 vector containing full-length SARS-CoV-2 S gene under control of the CMV promoter and full-length SARS-CoV-2 N gene under control of the human betaactin promoter.
c. rAd-S1-N: rAd5 vector using a fusion sequence combining the S1 region of SARS-CoV-2 S gene (including the native furin site between S1 and S2) with the full-length SARS-CoV-2 N gene.
d. rAd-S(fixed)-N: rAd5 vector containing a stabilized S gene with the transmembrane region removed under the control of the CMV promoter and full-length SARS-CoV-2 N gene under control of the human beta-actin promoter. The S gene is stabilized through the following modifications:

a. Arginine residues at aa positions 682, 683, 685 were deleted to remove the native furin cleavage site
b. Two stabilizing mutations were introduced: K986P and V987P
c. Transmembrane region was removed following P1213 and replaced with bacteriophage T4 fibritin trimerization foldon domain sequence^51^ (GYIPEAPRDGQAYVRKDGEWVLLSTFL)

All vaccines were grown in the Expi293F suspension cell-line (Thermo Fisher Scientific), purified by CsCl density centrifugation and provided in a liquid form for animal experiments.

### Animal Experiments

Studies were approved for ethics by the Animal Care and Use Committees (IACUC). All of the procedures were carried out in accordance with local, state and federal guidelines and regulations. Female 6-8 week old Balb/c mice were purchased from Jackson labs (Bar Harbor, ME). Because mice do not swallow pills, liquid formulations were instilled intranasally in 10 μl per nostril, 20 μl per mouse in order to test immunogenicity of the various constructs. Serum was acquired by cheek puncture at various timepoints.

### Antibody Assessment

#### ELISAs

Specific antibody titers to proteins were measured similarly to methods described previously^52^. Briefly, microtiter plates (MaxiSorp: Nunc) were coated in 1 carbonate buffer (0.1 M at pH 9.6) with 1.0 ug/ml S1 protein (GenScript). The plates were incubated overnight at 4°C in a humidified chamber and then blocked in PBS plus 0.05% Tween 20 (PBST) plus 1% BSA solution for 1 h before washing. Plasma samples were serially diluted in PBST. After a 2-h incubation, the plates were washed with PBST at least 5 times. Antibodies were then added as a mixture of anti-mouse IgG1-horseradish peroxidase (HRP) and anti-mouse IgG2a-HRP (Bethyl Laboratories, Montgomery, TX). Each secondary antibody was used at a 1:5,000 dilution. The plates were washed at least 5 times after a 1-h incubation. Antigen-specific mouse antibodies were detected with 3,3=,5,5=-tetramethyl-benzidine (TMB) substrate (Rockland, Gilbertsville, PA) and H2SO4 was used as a stop solution. The plates were read at 450 nm on a Spectra Max M2 Microplate Reader. Average antibody titers were reported as the reciprocal dilution giving an absorbance value greater than the average background plus 2 standard deviations, unless otherwise stated.

#### Antibody binding antibodies

To measure responses to both S1 and S2 simultaneously, A MULTI-SPOT^®^ 96-well, 2-Spot Plate (Mesoscale Devices; MSD) was coated with SARS CoV-2 antigens. Proteins were commercially acquired from a source (Native Antigen Company) that produced them in mammalian cells (293 cells). These were biotinylated and adhered to their respective spots by their individual U-PLEX linkers. To measure IgG antibodies, plates were blocked with MSD Blocker B for 1 hour with shaking, then washed three times prior to the addition of samples, diluted 1:4000. After incubation for 2 hours with shaking, the plates were washed three times. The plates were then incubated for 1 hour with the detection antibody at 1 μg/mL (MSD SULFO-TAG™ Anti-mouse IgG). After washing 3 times, the Read Buffer was added and the plates were read on the Meso QuickPlex SQ 120.

### SARS-CoV-2 Neutralization Assays

Neutralizing antibodies were routinely detected based on the SARS-CoV-2 Surrogate Virus Neutralization Test (sVNT) kit (GenScript). This ELISA-based kit detects antibodies that hinder the interaction between the receptor binding domain (RBD) of the SARS-CoV-2 spike glycoprotein and the ACE2 receptor on host cells, and is highly correlated to conventional virus neutralizing titers for SARS-CoV-2 infection of Vero cells^53^. The advantage of this approach is that the assay can be done in a BSL-2 laboratory. Sera from mice immunized with the candidate vaccines was diluted at 1:20, 1:100, 1:300, 1:500, 1:750 and 1:1000 using the provided sample dilution buffer. Sera from non-immunized mice was diluted 1:20. Lung samples were diluted 1:5, 1:20, and 1:100. Positive and negative controls were prepared at a 1:9 volume ratio following the provided protocol. After dilution, sera or lung samples were individually incubated at a 1:1 ratio with HRP-RBD solution for 30 minutes at 37°C. Following incubation, 100μl of the each HRP-RBD and sample or control mixture was added to the corresponding wells in the hACE2-precoated capture plate and once again incubated at 37°C for 15 minutes. Afterwards, wells were thoroughly washed and 100μl of the provided TMB (3,3=,5,5=-tetramethyl-benzidine) solution was added to each well and left to incubate for 15 minutes at room temperature (20-25°C). Lastly 50μl of Stop Solution was added to each well, and the plate was read on a Spectra Max M2 Microplate Reader at 450 nm. The absorbance of a given sample is inversely related on the titer of anti-SARS-CoV-2 RBD neutralizing antibody in a given sample. Per test kit protocol, a cut-off of 20% inhibition when comparing the OD of the sample versus the OD of the negative control was determined to be positive for the presence of neutralizing antibodies. Samples that were negative at the lowest dilution were set equal to ½ of the lowest dilution tested, either 10 for sera or 2.5 for lung samples.

Additional neutralizing antibodies responses were measured in some studies using a cVNT assay at Visimederi under BSL3 conditions. The cVNT assay has a readout of Cytopathic Effect (CPE) to detect specific neutralizing antibodies against live SARS-COV-2 in animal or human samples. The cVNT/CPE assay permits the virus to makes multiples cycles of infection and release from cells; its exponential grow in few days (usually 72 hours of incubation) causes the partial or complete cell monolayer detachment from the surface of the support, clearly identifiable as CPE. Serum samples are heat inactivated for 30 min at 56°; two-fold dilutions, starting from 1:10 are performed then mixed with an equal volume of viral solution containing 100 TCID_50_ of SARS-CoV-2. The serum-virus mixture is incubated for 1 hour at 37° in humidified atmosphere with 5% CO2. After incubation, 100 μL of the mixture at each dilution are added in duplicate to a cell plate containing a semiconfluent Vero E6 monolayer. After 72 hours of incubation the plates are inspected by an inverted optical microscope. The highest serum dilution that protect more than 50% of cells from CPE is taken as the neutralization titer.

#### Lung IgA ELISAs

Two weeks after the final immunization (day 28 of the study), mice were sacrificed and bled via cardiac puncture. Lungs were removed and snap frozen at −80 °C. On thawing, lungs were weighed. Lungs were homogenized in 150 μl DPBS using pellet pestles (Sigma Z359947). Homogenates were centrifuged at 1300rpm for 3 minutes and supernatants were frozen. The total protein content in lung homogenate was evaluated using a Bradford assay to ensure equivalent amounts of tissue in all samples before evaluation of IgA content. Antigen-specific IgA titers in lungs were detected using a mouse IgA ELISA kit (Mabtech) and pNPP substrate (Mabtech). Briefly, MaxiSorp plates (Nunc) were coated with S1 or S2 (The Native Antigen Company; 50 ng/well) in PBS for overnight adsorption at 4°C and then blocked in PBS plus 0.05% Tween 20 (PBST) plus 0.1% BSA (PBS/T/B) solution for 1 h before washing. Lung homogenates were serially diluted in PBS/T/B, starting at a 1:30 dilution. After 2 hours incubation and washing, bound IgA was detected using MT39A-ALP conjugated antibody (1:1000), according to the manufacturer’s protocol. Plates were read at 415nm. Endpoint titers were taken as the x-axis intercept of the dilution curve at an absorbance value 3x standard deviations greater than the absorbance for naïve mouse serum. Non-responding animals were set a titer of 15 or ½ the value of the lowest dilution tested.

### T cell Responses

Spleens were removed and placed in 5 ml Hanks Balanced Salt Solution (with 1M HEPES and 5% FBS) before pushing through a sterile strainer with a 5ml syringe. After RBC lysis (Ebiosolutions), resuspension, and counting, the cells were ready for analysis. Cells were cultured at 5e5 cells/well with two peptide pools representing the full-length S protein at 1 μg/ml (Genscript) overnight in order to stimulate the cells. The culture media consisted of RPMI media (Lonza) with 0.01M HEPES, 1X l-glutamine, 1X MEM basic amino acids, 1X penstrep, 10% FBS, and 5.5e-5 mole/l beta-mercaptoethanol. Antigen specific IFN-γ ELISPOTs were measured using a Mabtech kit. Flow cytometric analysis was performed using an Attune Flow cytometer and Flow Jo version 10.7.1, after staining with the appropriate antibodies. For flow cytometry, 2e6 splenocytes per well were incubated for 18 hours at 37°C with peptide pools representing full-length S at either 1 or 5 ug/ml, adding Brefeldin A (ThermoFisher) for the last 4 hours of incubation. The antibodies used were APC-H7 conjugated CD4, FITC conjugated CD8, BV650 conjugated CD3, PerCP-Cy5.5 conjugated IFN-y, BV421 conjugated IL-2, PE-Cy7 conjugated TNFa, APC conjugated IL-4, Alexa Fluor conjugated CD44, and PE conjugated CD62L (BD biosciences).

## ACKNOWLEDGEMENTS

The authors would like to thank Alessandro Torelli, Laura Palladino, Emanuele Montomoli and the team at VisMederi for running neutralizing antibody titers (BSL3) and Susan Johnson for critically reviewing the manuscript.

## AUTHORSHIP

ACM, MC, and SNT designed the experiments; ACM and SNT wrote the manuscript; NP, KPT, and KL performed immunological assays; ACM, NP, KPT, KL, MC, and SNT analyzed the data. EGD and SNT designed the vaccine vectors.

