## Supplemental Table 1 for "Pre-clinical studies of a recombinant adenoviral mucosal vaccine to prevent SARS-CoV-2 infection"

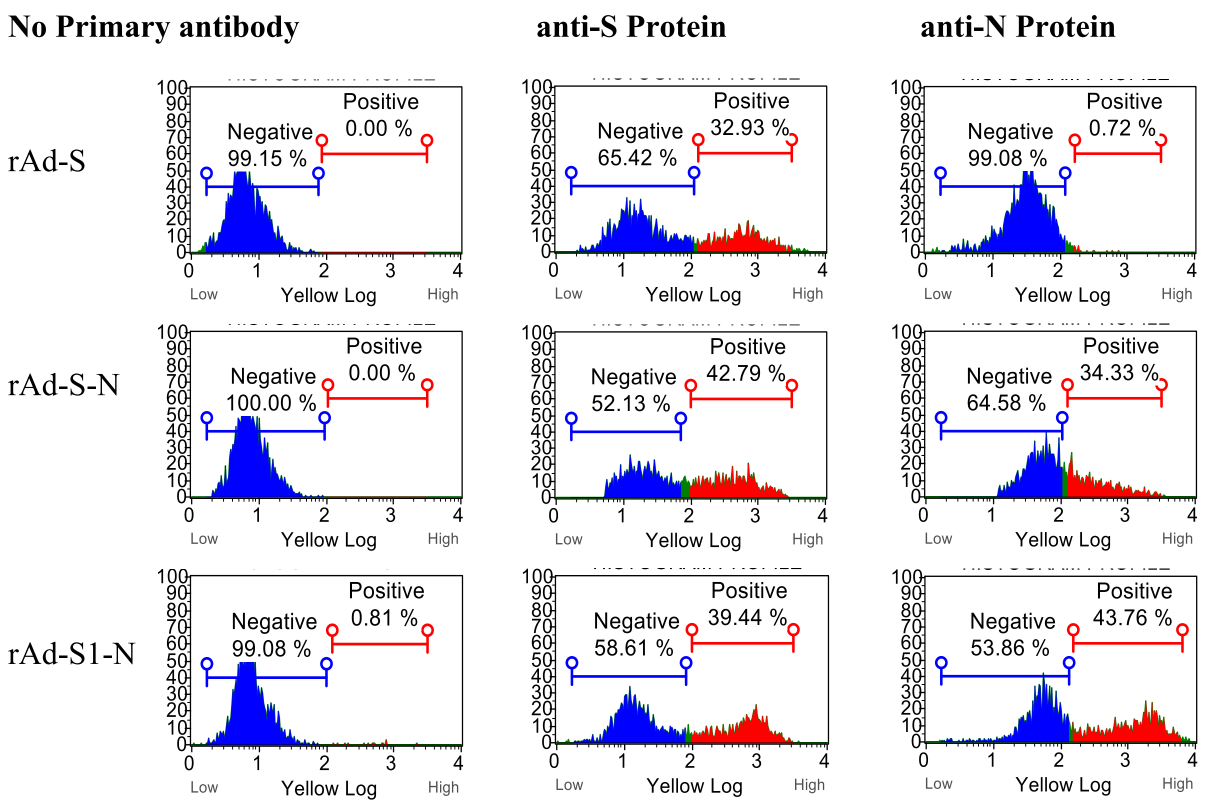


*Figure 1. Expression of the antigens in human cells post infection. 293 cells were transduced with one of the recombinant adenoviral vectors (rAd-S, rAd-S-N, rAd-S1-N) and stained for expression of either the spike (S) or nucleocapsid (N) protein. The appropriate virus made the correct proteins.*
